# Rhythms of early life: gut microbiota rhythmicity and circadian maturation in infants

**DOI:** 10.1101/2025.01.06.631487

**Authors:** Christophe Mühlematter, Dennis S. Nielsen, Josue L. Castro-Mejía, Jean-Claude Walser, Sarah F. Schoch, Salome Kurth

## Abstract

**Background:** The human gut microbiota undergoes daily fluctuations, yet its interaction with sleep-wake patterns during infancy remains largely uncharted. This study aims to elucidate the relationship between gut microbiota rhythmicity and the development of sleep patterns in infants over the first year of life. We continuously monitored 162 healthy infants across multiple days at 3, 6, and 12 months of age using ankle actigraphy and 24-hour diaries. The Circadian Function Index (CFI) was computed as a proxy for sleep-wake rhythm maturation. Stool samples were collected to profile gut microbiota taxa composition via 16S rRNA gene amplicon sequencing, and microbial oscillations were assessed through sine and cosine fitting to detect 24-hour patterns.

**Results:** Our findings revealed that the relative abundance of bacterial taxa exhibited rhythmic patterns, with 26 zOTUs (1.74%) following a sine pattern and 100 zOTUs (6.69%) displaying cosine rhythmicity. Cosine rhythmicity became more pronounced with age, showing strong maturation: 7 zOTUs at 3 months, 2 zOTUs at 6 months, and 86 zOTUs at 12 months.

Notably, 105 zOTUs (7.02%) were associated with CFI, demonstrating a significant relationship between gut microbiota rhythms and sleep development. Among these, 27 zOTUs with sine dynamics and 96 zOTUs with cosine dynamics were linked to CFI, with this association strengthening as infants aged.

**Conclusions:** These results highlight the increasing synchronization between gut microbiota rhythmicity and sleep-wake cycles during infancy, pointing to a critical window for potential health interventions. This novel observation, previously reported in rodents and adults, underscores the role of gut microbiota in early human development, offering new avenues for enhancing developmental outcomes through targeted interventions.

## Introduction

Infancy represents a critical developmental phase of the human brain. During early childhood sleep undergoes pronounced maturation, a process essential for cognitive and neurophysiological development (Timofeev et al., 2020). Core electrophysiological features of sleep, such as slow waves and spindles, have been identified as biomarkers for the maturation of neuronal connectivity and function (Jaramillo et al., 2023; Kurth et al., 2012).

Sleep furthermore affects immuno-regulation (Besedovsky et al., 2019), cardiovascular function (Korostovtseva et al., 2021), and metabolic balance (Stich et al., 2022), underscoring the necessity of adequate sleep to both neurological development and physiological health. Early human life is characterized by notable variability in individual sleep behavior (Iglowstein et al., 2003; Schoch, Huber, et al., 2020). Despite the substantial health implications, such as effects on cognitive development and obesity risk (Li et al., 2022; Morales-Muñoz et al., 2021), the underlying drivers of this variability remain largely obscure. The complexity of capturing sleep maturation through continuous monitoring presents a substantial challenge, hindering a thorough understanding of infant sleep patterns. During infancy, neurophysiology and behavior are highly modifiable, allowing for effective re-shaping of developmental trajectories through timely interventions (Sinthong & Ngernlangtawee, 2024). For example, early sleep interventions have been shown to extend nighttime sleep duration, reduce night wakings, and promote self-soothing behaviors, all of which contribute to improved sleep quality and support healthy development. This highlights a promising approach to enhancing sleep regulation and fostering positive developmental outcomes.

Given the multifaceted factors influencing sleep, recent research has uncovered a bidirectional relationship between sleep and gut microbiota—the microorganisms inhabiting the gastrointestinal tract (Sen et al., 2021). Modifying the gut microbiota through antibiotics can disrupt sleep patterns, such as reducing non-rapid eye movement sleep (NREMS) and increasing REM sleep episodes in mice, or decreasing slow-wave sleep (SWS) in humans and rats, suggesting a role for microbiota in maintaining healthy sleep architecture (Brown et al., 1990; Nonaka et al., 1983; Ogawa et al., 2020). Conversely, sleep disruptions have been found to alter gut microbiota composition (Bowers et al., 2020; Thaiss et al., 2014). In adults, a fraction of variability in sleep quality and duration is explained by gut microbiota profiles, underscoring the interconnectedness of sleep to microbial signatures (Grosicki et al., 2020; Smith et al., 2019). The sleep-gut connection is not limited to adulthood, but is observed in adolescence, childhood, and infancy; likely it primes health outcomes across the lifespan (Carpena et al., 2024; Schoch et al., 2022; Wang et al., 2022; Xiang et al., 2023).

Similar to sleep, variability in individual gut microbiota during infancy is large (Hill et al., 2017), and defines critical health aspects such as metabolism (Altaha et al., 2022), immune function (Lubin et al., 2023), and neurodevelopment (Sordillo et al., 2019). Colonization of the infant gut begins at birth, and across the first three years of life microbial diversity, stability, and complexity increase. The succession of microbial communities in a healthy infant gut is influenced by environmental and dietary factors, such as mode of birth, antibiotic usage, and dietary habits (Catassi et al., 2024; Mercer et al., 2024; Pantazi et al., 2023).

Furthermore, gut microbiota have recently been found to undergo “diurnal” dynamics, demonstrating variance in abundance and function across the day (Thaiss et al., 2014; Voigt et al., 2014). Animal models show that diurnal microbial dynamics are tied to the host’s circadian rhythm, as bacterial diurnal variation is partially regulated by their host’s clock genes (Per1/2, Bmal1) (Heddes et al., 2022; Thaiss et al., 2016). However, this interaction between the host’s circadian rhythm and bacterial rhythm appears to be bidirectional: in germ-free mice, the absence of microorganisms disrupts the intestinal epithelial cell clock, decreasing the expression of core clock genes such as Bmal1 and Cry1, while increasing Per1 and Per2. This disruption alters metabolic homeostasis and rhythmic expression in the intestinal cells (Mukherji et al., 2013). Further evidence supporting the role of the gut microbiome as a “bottom-up” regulator of the circadian clock is provided by observations that microbial metabolites modulate the peripheral clock in mice (Tahara et al., 2018). This suggests that the interaction between the gut microbiota and the host’s biological clock extends beyond mere temporal coordination but also includes but also includes regulatory effects on the host’s circadian clock function.

In addition to these observations in adult animals, novel studies suggest that diurnal variations also occur in the gut microbiota of infants (Heppner et al., 2024). However, their connection to the infants’ circadian rhythm remains unknown. Addressing this phenomenon requires longitudinal data across infancy, collected from large, well-controlled cohorts with objective and continuous assessments of sleep. Ankle-actigraphy captures physiological acceleration, thereby providing reliable data that captures the large individual variability in infant rest-activity patterns (Schoch, Kurth, et al., 2020; Werner et al., 2008). Applying the Circadian Function Index (CFI) (Ortiz-Tudela et al., 2010) can serve as a proxy for circadian maturation; facilitating the investigation of its relationship with gut microbiota dynamics.

We hypothesized that individual gut microbiota profiles are associated with the maturation of infant circadian rhythm, using an increased CFI as proxy for this maturation. This relationship would indicate that as the circadian rhythm matures, distinct shifts in gut microbiota composition can be observed. Additionally, we predicted a maturation of diurnal rhythm in gut microbiota composition across the first year of life. Specifically, we anticipated that the proportion of rhythmic zOTUs would increase with age. Third, we expected that the diurnal microbial rhythm is linked to infants’ sleep regulation, quantified by the CFI. A linkage would indicate that as the circadian rhythm develops and strengthens, the rhythmic expression of the gut microbiota also becomes more pronounced.

## Methods

### Study Population

This study was conducted in Switzerland and included 162 healthy infants assessed at ages 3-12 months from 2016 to 2019 (Schoch et al., 2022). Infants included in the study were term-born, showed no signs of acute or chronic illness, were delivered vaginally, and received at least 50% of their nutrition from breast milk at the time of enrollment. Exclusion criteria encompassed the use of antibiotics or sleep-affecting medications, chronic illnesses, or disorders of the central nervous system. The cantonal ethics committee of Zurich, Switzerland, approved the project (BASEC 2016-00730), in line with the principles of the Helsinki Declaration. Before study enrollment, parents received detailed explanations about the study procedures before providing written consent.

### Study design

Families and infants were longitudinal assessed at 3, 6, and 12 months of age. For each assessment, actigraphy data was collected for 11 consecutive days (9.16 ± 1.23 days) using a GENEactiv sensor (Activinsights LtD, Kimbolton, UK, 43×40×13mm, MEMS sensor, 16g, 30 Hz frequency) attached to the infant’s left ankle (either with a paper strap or in a modified sock). Concurrently, caregivers completed a 24h paper-pencil sleep diary to document sleep and wake periods, any temporary removal of the device, and external movements during sleep (*e.g.*, such as when the infant was in a moving vehicle) (Schoch et al., 2019; Werner et al., 2008). Additionally, caregivers collected at least one infant stool sample during each assessment, for which clock time was documented. Demographic data and complementary information was collected with online questionnaires at each assessment timepoint (SoSci Survey; (Leiner, 2021).

### Data processing

Actimetric activity counts were processed in Matlab (R2016b), by using binary files extracted through GENEactiv PC Software (version 3.1). Transformation involved applying a 3–11 Hz bandpass filter and compressing the signal into 15-second bins. Acceleration data from all three axes was combined using a sum of squares. To enhance data accuracy, actigraphy output was integrated with information provided by parents in the 24-hour diaries, particularly regarding instances when the actimeter was not worn, instances of external movement during sleep (e.g., while in a car), as well as cases of poor fit between actimetry data and diary records. The acceleration signal was compressed to one data point per minute through data summation (Schoch, Huber, et al., 2020). This compressed acceleration data was then used to compute the CFI as a proxy for circadian maturation.

### Circadian Function Index

While previous studies used infant ankle actimetry to assess sleep quantity and continuity (Schoch, Huber, et al., 2020; Schoch, Kurth, et al., 2020), the use of CFI captures the maturation of sleep-wake rhythmicity across several days, introducing a novel perspective for pediatric circadian development. CFI measures the regularity of activity-rest patterns with a score from 0 to 1 (Ortiz-Tudela et al., 2010). The computation of CFI relies on 3 measures, computed through the R package “nparact” version 0.8 (Blume et al., 2016): Intradaily Variability (IV), Interdaily Stability (IS), and Relative Amplitude (RA). IV determines the frequency and extent of transitions between rest and activity within a 24-hour day, such that a lower IV represents more consolidated rest-activity periods, whereas a higher IV suggests frequent transitions and thus fragmented sleep patterns. IS quantifies the similarity of activity patterns across days. A higher IS represents a more stable day-to-day activity-rest pattern.

RA measures the amplitude between the most active and least active periods within a 24-hour cycle. A higher RA indicates a more pronounced difference between peak activity and rest periods.

For each infant, these three scores were averaged across days of continuous actigraphy data collection, covering an average period of 9.16 ± 1.23 days across all ages (3, 6, and 12 months). Specifically, at 3 months, infants were assessed over an average of 9.28 ± 1.17 days. At 6 months, the average was 9.14 ± 1.46 days. By 12 months, the average assessment period was 9.04 ± 0.99 days. The CFI was then calculated with the following formula:

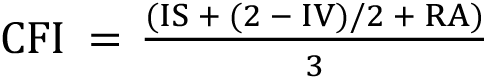

This formula combines each component to provide an overall measure of circadian maturation, with higher CFI values reflecting more regular and stable activity-rest patterns.

### DNA extraction from stool and processing of gut microbiota

Gut microbiota profiles were computed from stool samples collected at infant ages 3, 6, and 12 months. Upon collection, samples were stored in the participants’ fridge (approximately 4°C) before being transported at a consistent temperature within 96 hours to the laboratory. Upon arrival, samples were divided into 200 mg aliquots and preserved at-50 °C until processing. Total DNA extraction was performed using a PowerSoil DNA Isolation Kit (MOBIO Laboratories, Carlsbad, CA, USA), following the manufacturer’s guidelines with small modifications (for details see (Schoch et al., 2022). 16S rRNA gene amplicon libraries were prepared using specific primers targeting the V3 region (Ovreås et al., 1997), equipped with adapters compatible with the Nextera Index Kit® (Illumina, CA, USA): NXt_338_F: 5′-TCG TCG GCA GCG TCA GAT GTG TAT AAG AGA CAG ACW CCT ACG GGW GGC AGC AG-3′ and NXt_518_R: 5′-GTC TCG TGG GCT CGG AGA TGT GTA TAA GAG ACA GAT TAC CGC GGC TGC TGG-3′. Subsequent steps for amplification profiling, barcoding, purification of amplicon libraries, and sequencing were carried out as previously described (Krych et al., 2018). Initial quality assessment of the raw datasets (2×151bp, paired-end) was conducted using FastQC58. Overlapping reverse and forward reads were trimmed using SeqKit59, and reads were merged with FLASH (v1.2.11) (Magoč & Salzberg, 2011), allowing for overlaps between 15 and 300, with a maximum mismatch density of 0.25. Cutadapt (v1.12) (Martin, 2011) was utilized for primer region trimming, maintaining an error rate below 0.01. Prinseq (Schmieder & Edwards, 2011) was used for quality filtering by discarding reads containing ambiguous nucleotides or with an average quality score below 20. Zero-radius Operational Taxonomic Units (zOTUs) were identified using usearch (UNOISE3 v10.0.240) (Edgar & Flyvbjerg, 2015), and taxonomic classifications were determined based on the Greengenes database (DeSantis et al., 2006).

To mitigate batch effects during stool sample processing, batch run was incorporated as a control variable in all statistical models. Samples with fewer than 50,000 reads were excluded from the study (n=6). The remaining samples were rarefied to the lowest observed read count of 50,268. This process resulted in 1430 amplicon sequencing variants.

Autoclaved water samples, subjected to the same experimental procedures, served as negative controls and exhibited read counts ranging from 2 to 150,785. zOTU2 (Enterobacteriaceae), identified in 30% of the negative controls and 10% of the stool samples, showed no significant impact on the studied gut microbiota markers. Only bacterial taxa present in at least 20% of the samples from all collection points (3-, 6-, and 12-months) and accounting for at least 1% of total bacterial counts were included in subsequent analyses. This selection process resulted in 1451 zOTUs included in the analysis.

### Rhythmicity Analysis

To assess diurnal rhythmicity in the relative abundance of specific bacterial taxa (zOTUs), we adapted a cosinor analysis approach. Cosinor analysis utilizes sinusoidal models to capture periodicity in time-series data and is thus particularly well-suited for biological rhythms (Cornelissen, 2014). We transformed the clock times of stool sample collection into numerical values representing the time of day, and then applied sine and cosine transformations to these values. By fitting sine and cosine functions, we could evaluate whether zOTU relative abundance exhibited patterns conforming to a 24-hour cycle. This method allowed us to detect and quantify cyclical variations in the relative abundance of microbial populations.

### Statistical Analysis

Data processing, modeling, visualization, and statistical analysis was performed using R version 4.2.1 and packages ggplot2 (Wickham, 2016), phyloseq (McMurdie & Holmes, 2013), and lme4 (Bates et al., 2015). We applied linear mixed-effects models to compute associations between the relative abundance of zOTUs (dependent variable) and CFI, clock time of stool sampling (fitted to sine and cosine functions), and age group, allowing us to examine microbial rhythmicity. Microbial rhythmicity was assessed by modeling sine and cosine terms separately for each zOTU, and a zOTU was considered rhythmic if either the sine or cosine term showed a significant association (FDR-corrected p-value < 0.05) with relative abundance at a given age. We further incorporated interactions between sine and cosine fits with CFI to assess the combined effects of infant circadian rhythm and microbial zOTU rhythmicity. For rhythmic zOTUs, we reported the relative proportions of phyla. Control variables in the models included sex (0 for male, 1 for female), breastfeeding status (0 for not or rarely breastfed, 1 for occasionally, regularly, or daily breastfeeding) and sequencing run (categorical, with levels 1 to 5). Random intercepts for each participant were included to capture individual-level variability. The alpha level was set to P < 0.05, and to address multiple comparison concerns, we applied False Discovery Rate (FDR) correction to the p-values.

## Results

Key demographic variables including antibiotic use, feeding method (percentage of breastmilk, formula, solids), and age at solid food introduction were summarized for each assessment point (3, 6, and 12 months, Table 1).

**Table 1.**
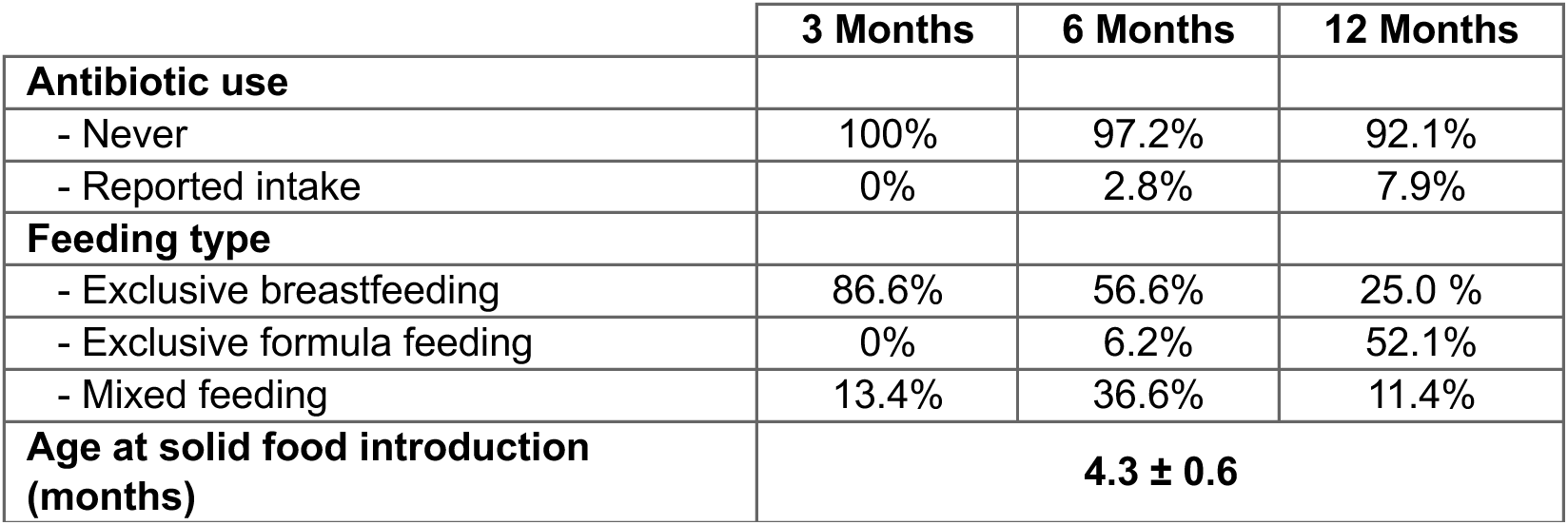
Participant demographics at 3, 6, and 12 months, including antibiotic use, feeding type, and age at solid food introduction.

|  | 3 Months | 6 Months | 12 Months |
| --- | --- | --- | --- |
| **Antibiotic use** |  |  |  |
| - Never | 100% | 97.2% | 92.1% |
| - Reported intake | 0% | 2.8% | 7.9% |
| **Feeding type** |  |  |  |
| - Exclusive breastfeeding | 86.6% | 56.6% | 25.0 % |
| - Exclusive formula feeding | 0% | 6.2% | 52.1% |
| - Mixed feeding | 13.4% | 36.6% | 11.4% |
| **Age at solid food introduction (months)** | **4.3 ± 0.6** |  |  |

To assess whether stool sampling times were consistent across participants and age groups, we tested for potential differences in the timing of stool collection across timepoints (Figure 1), which was not the case (Kruskal-Wallis test, *p* = 0.75; all pairwise Wilcoxon tests *p* = 1 after Bonferroni correction).

**Figure 1.**
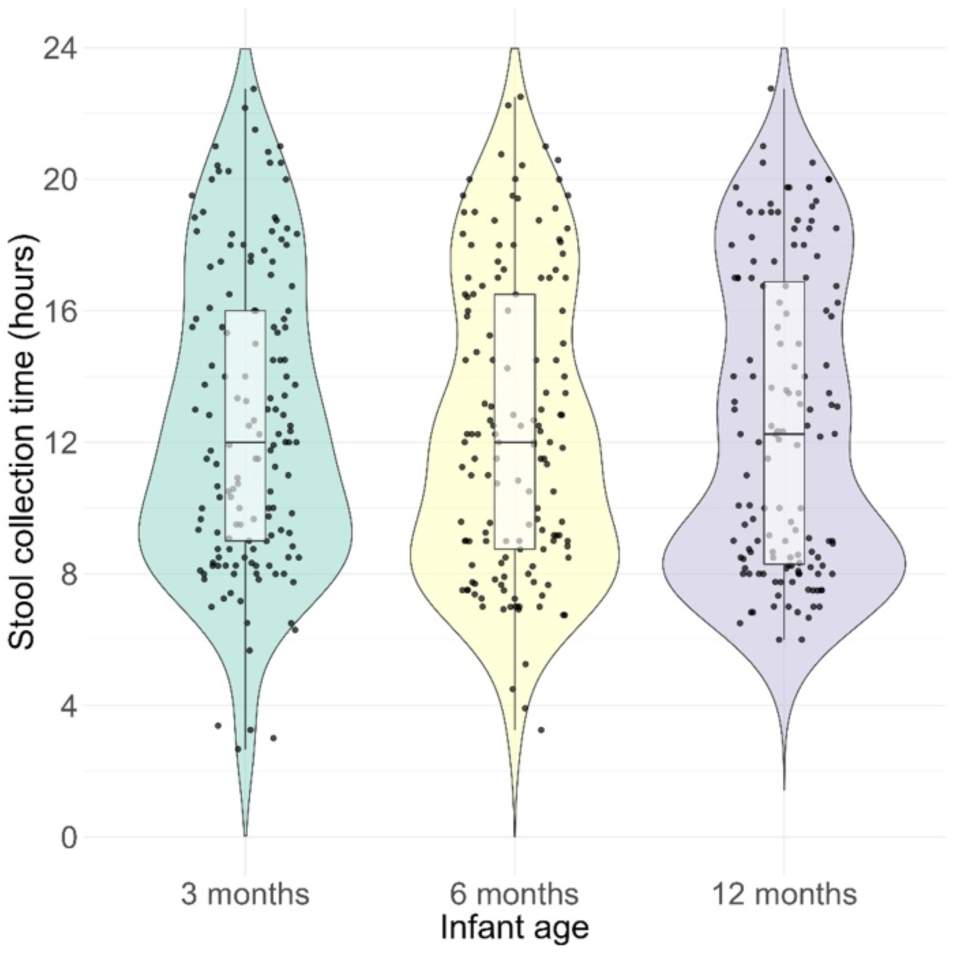
Timing of stool sample collection at each timepoint. Violin and scatter plots depict the distribution of collection times (in decimal hours) at 3, 6, and 12 months. Each dot represents one sample. No significant differences were observed between timepoints (Kruskal-Wallis test, *p* = 0.75).

### Circadian maturation across infancy

Across the first year of life, CFI increased with infant age, with mean CFI values of 0.59 ± 0.07 at 3 months, 0.64 ± 0.06 at 6 months, and 0.70 ± 0.05 at 12 months (Figure 2). CFI increased significantly from 3 months to 6 months (p < 0.001, 95% C.I. = 0.04, 0.08; Tukey post-hoc tests), from 3 months to 12 months (p < 0.001, 95% C.I. = 0.10, 0.13), and from 6 months to 12 months (p < 0.001, 95% C.I. = 0.042, 0.07).

**Figure 2.**
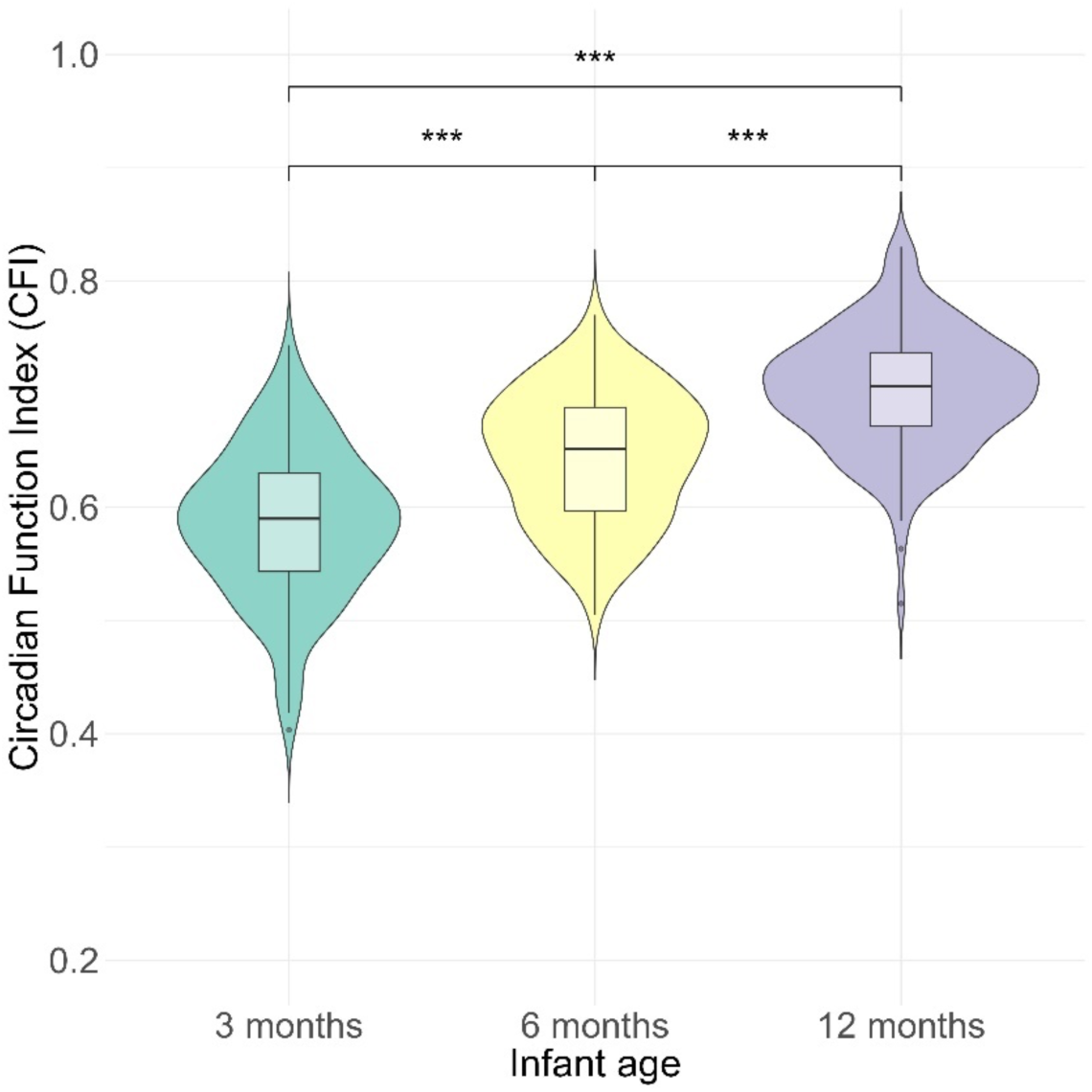
Age-related increase in sleep regulation across infancy assessed with Circadian Function Index (CFI) (Tukey post-hoc tests, *** p < 0.001). Central horizontal lines refer to median, the box spans the interquartile range (IQR), covering the middle 50% of data from the 25th to the 75th percentile. Whiskers extend to the highest and lowest values within 1.5 times the IQR from box edges. Points beyond the whiskers represent outliers.

### Association between circadian maturation and gut microbial taxa

To define whether infant CFI underlies differences in gut microbial taxonomic abundance, we analyzed associations between zOTUs relative abundance and CFI (rhythmicity of zOTU abundance is presented subsequently). This identifies the bacterial taxa linked to the infants’ circadian maturational status. We observed both positive and negative correlations between zOTUs and CFI. A total of 105 zOTUs showed significant associations with CFI, representing 7.02% of all identified zOTUs across samples. The strength of this association increased with age: at age 3 months, only 10 zOTUs were linked to CFI, increasing to 17 zOTUs at 6 months, and 87 zOTUs at 12 months. Thus, over the first year of life, a progressively stronger association between infant gut microbiota and sleep rhythm emerges.

Among all 105 zOTUs associated with CFI, 74 showed positive correlations (Figure 3A), indicating that infants with higher CFI had increased relative abundance of these bacterial taxa. At 3 months, among the 3 zOTUs positively correlated with the CFI were 1 Actinobacteria, 1 Bacteroidetes and 1 from unknown phylum (Figure 3B). At 6 months *Actinobacteria* was the primary phylum associated with CFI, while at 12 months it was mainly *Actinobacteria* and *Firmicutes*.

**Figure 3.**
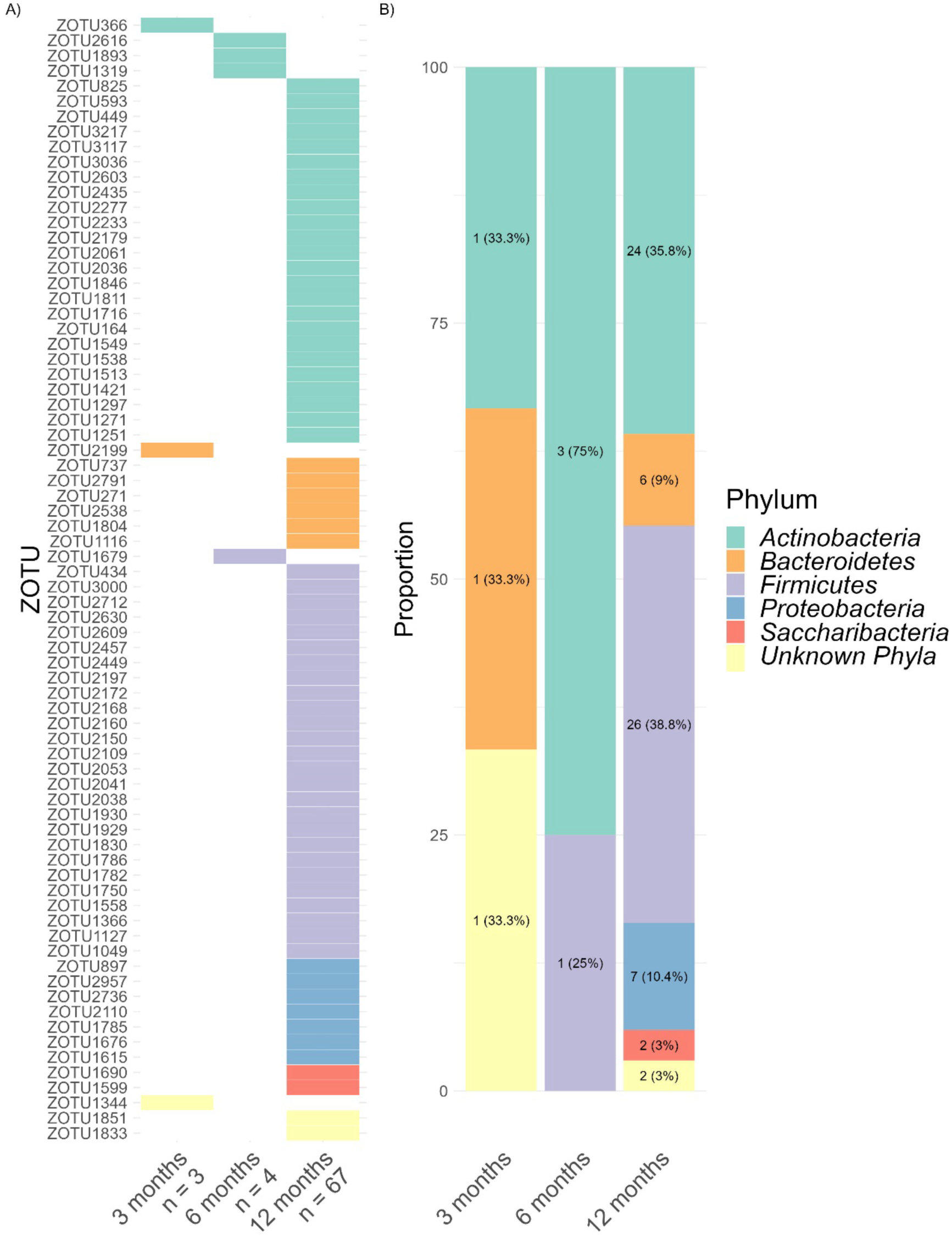
A) Heatmap of zOTUs (n = 74) with relative abundance positively associated with infant CFI (p < 0.05), specified for each age group. Values below age groups indicate the number of zOTUs. B) Distribution of phyla positively linked to infant CFI.

24 zOTUs were negatively linked with CFI, indicating that more mature child sleep regulation decreased the relative abundance in these zOTUs (Figure 4A). No negative association between CFI and gut microbiota relative abundance was observed at 3 months, while it increased to 6 zOTUs at 6 months and 18 zOTUs at 12 months. At 6 months, Actinobacteria was the most frequently associated phylum with CFI, while at 12 months it was Firmicutes (Figure 4B).

**Figure 4.**
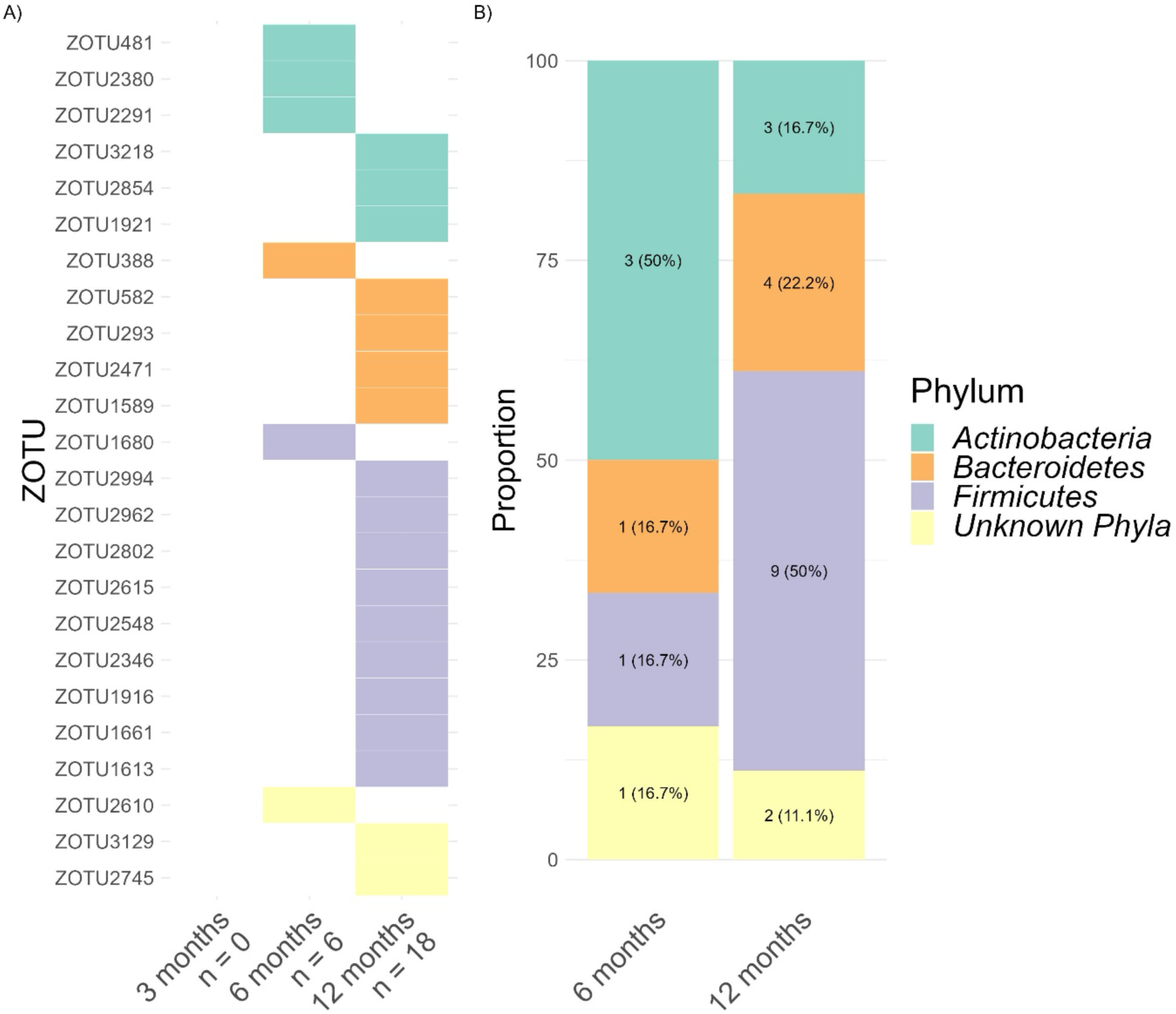
A) Heatmap of zOTUs (n = 24) with relative abundance negatively associated with infant CFI (p < 0.05), illustrated for each age group. Values below age groups represent the number of associated zOTUs. B) Distribution of zOTUs phyla negatively linked to infant CFI.

Interestingly, for 7 zOTUs, the direction of the association between zOTUs relative abundance and CFI was age dependent (Figure 5A). Specifically, at 3 months 6 zOTUs showed negative associations, which shifted to positive associations at 6 and/or 12 months. Conversely, one zOTU exhibited the opposite: showing positive correlation at 3 months shifting to negative at 6 and 12 months. Among these 7 zOTUs, at 3 and 6 months, there were 2 from the phylum *Firmicutes*, 1 from *Actinobacteria*, 1 from *Bacteroidetes*, 1 from *Proteobacteria* and 2 from unknown phyla. By 12 months, there was 1 zOTU from *Proteobacteria* and 1 from an unknown phylum (Figure 5B).

**Figure 5.**
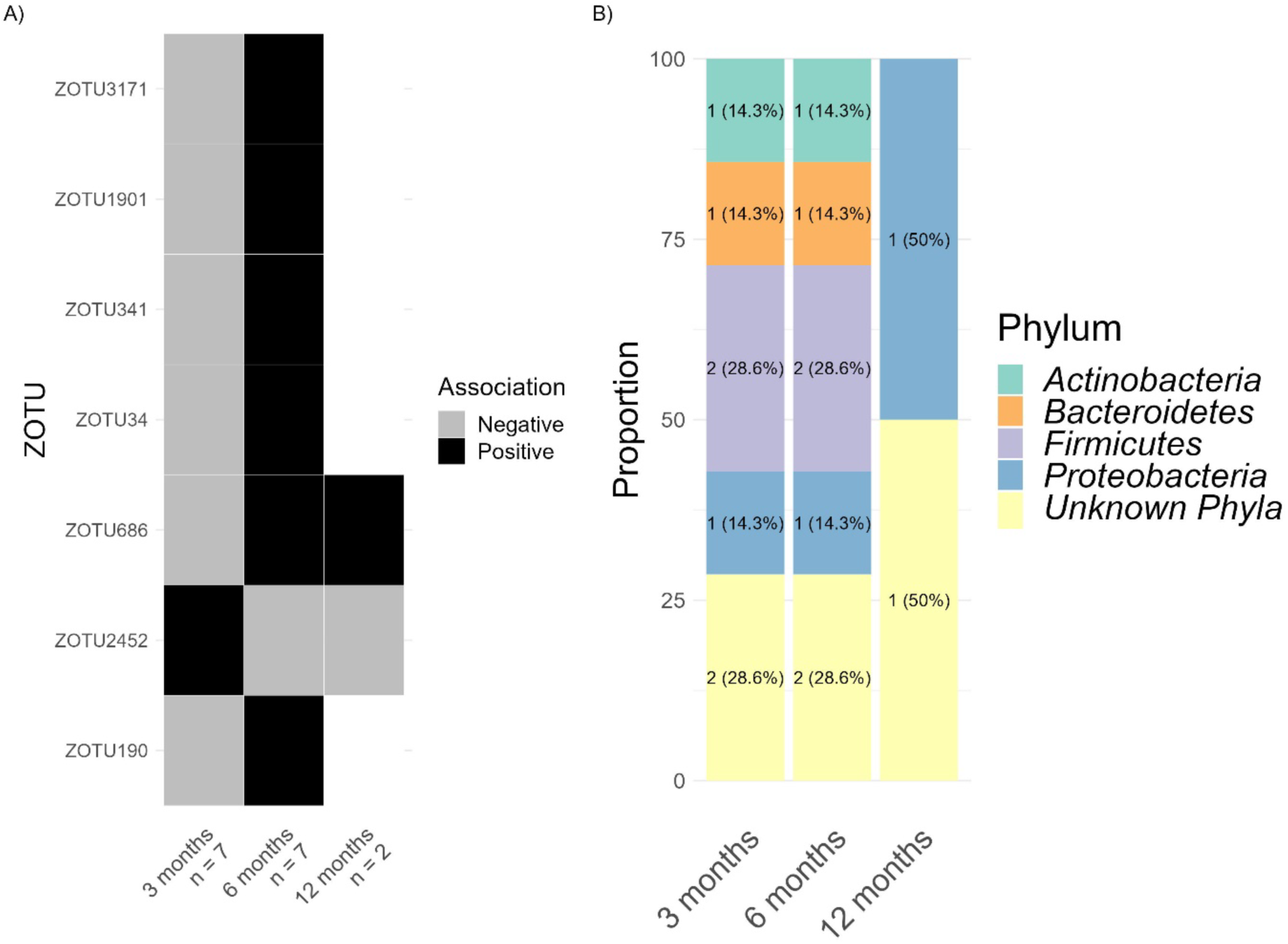
A) Heatmap of zOTUs (n = 7) with their relative abundance positively (black) or negatively (grey) associated with infant CFI (p <0.05). Values below age groups represent the number of zOTUs identified. B) Distribution of zOTU’s bacterial phyla linked to infant CFI, showing changes dependent on age.

### Rhythmic patterns in gut microbiota relative abundance

Subsequently, we investigated whether rhythmicity in gut microbial relative abundance was present during infancy, by considering the influence of clock time of stool sampling. The clock time of stool sampling was fitted to sine and cosine curves capturing potential diurnal effects.

Overall, the relative abundance of 26 zOTUs (i.e., 1.74% of total identified zOTUs) followed a sine pattern, indicating rhythmic diurnal variation in their abundance. The number of rhythmic zOTUs remained relatively constant across age (9 rhythmic zOTUs at 3 month, 10 rhythmic zOTUs at 6 months and 14 rhythmic zOTUs at 12 months; Figure 6A). *Actinobacteria* was the main rhythmic phyla undergoing a sine pattern at 3 months (33.3%) and 6 months (40%; Figure 6B). *Bacteroidetes* and *Firmicutes* represented rhythmic phyla at 12 months (each 42.9%).

**Figure 6.**
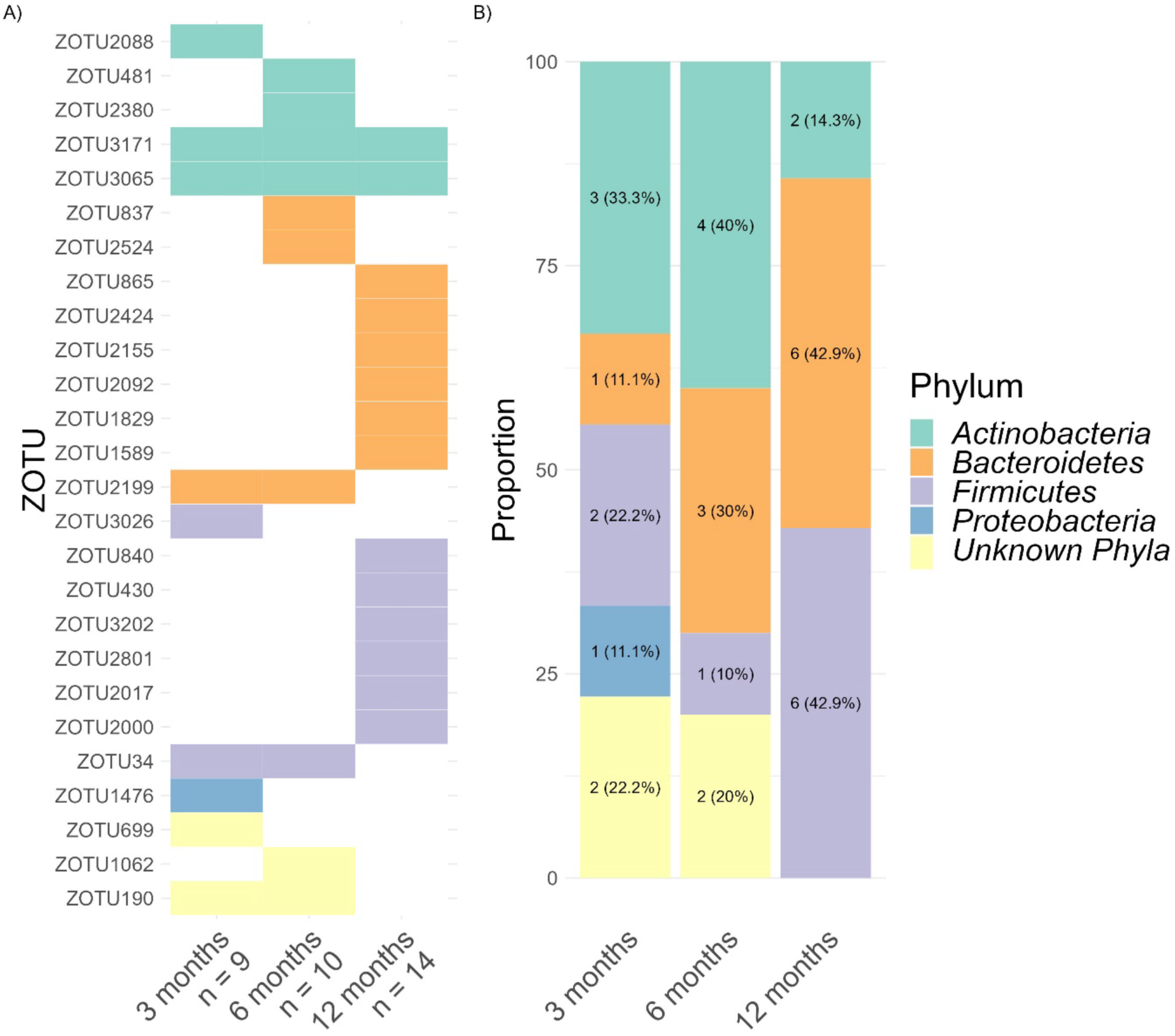
A) Heatmap of zOTUs (n = 26) with rhythmic relative abundance following a sine pattern (p < 0.05). Values below age groups represent the number of rhythmic zOTUs abundance. B) Distribution of zOTUs phyla following a sine rhythm.

Next, we applied the cosine approach to zOTU relative abundance. Compared to sine fitting, cosine fitting revealed that a greater number of zOTUs (100 in total, accounting to 6.69% of all zOTUs) exhibited rhythmic dynamics (Figure 7A). More pronounced than sine, rhythmic cosine dynamics unraveled an age-dependent increase, starting with 7 zOTUs at 3 months, increasing from 12 zOTUs at 6 months to 86 zOTUs at 12 months. The main cosine rhythmic zOTUs bacteria at 3 months were of unknown phylum, while for 6 and 12 months the majority were assigned to *Firmicutes* (respectively 41.7% and 45.3%, Figure 7B).

**Figure 7.**
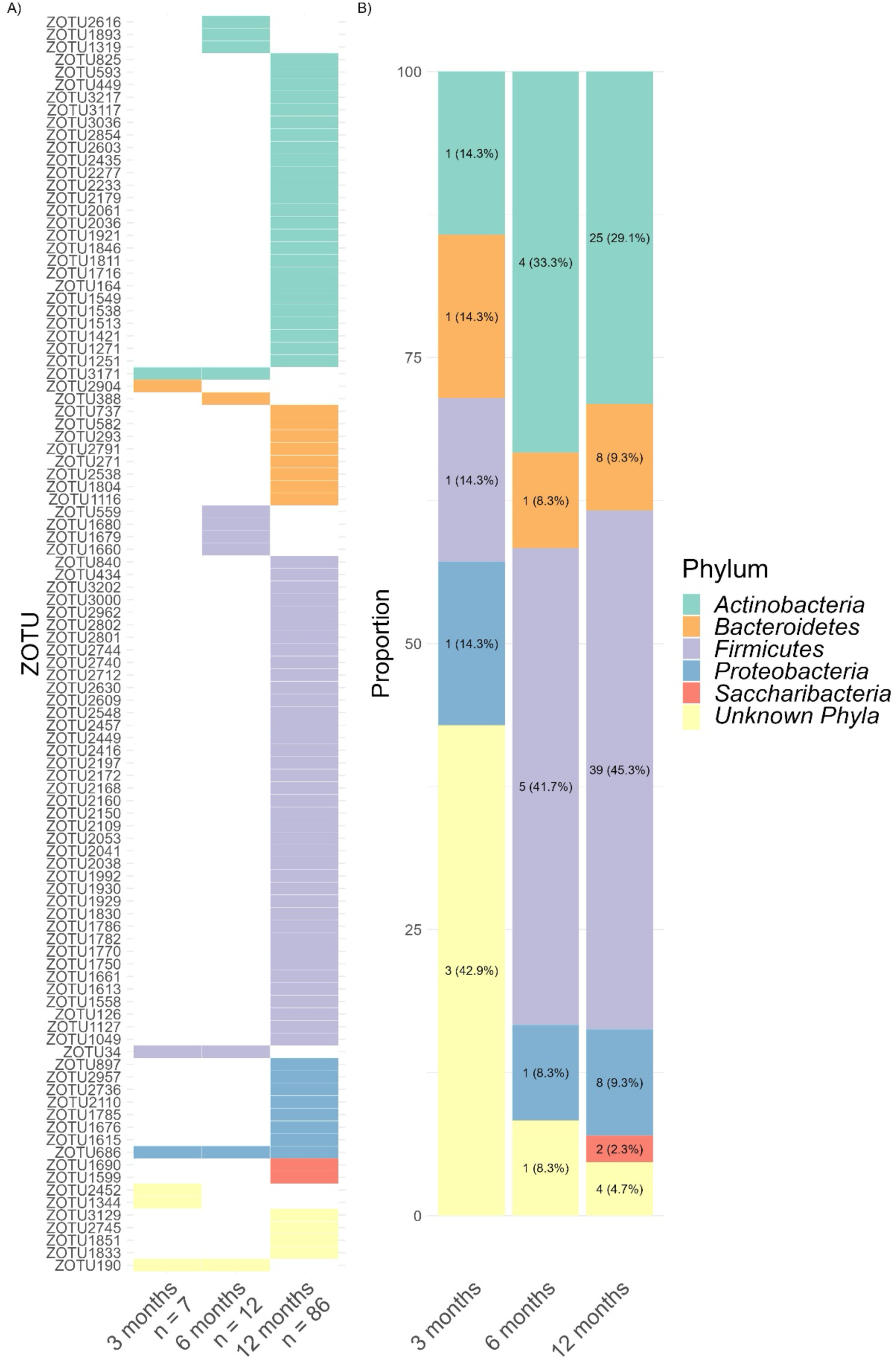
A) Heatmap of zOTUs (n = 100) whose relative abundance followed a cosine rhythm (p < 0.05). Values below age groups represent the number of zOTUs identified. B) Distribution of zOTU phyla expressed as a cosine pattern.

### Gut microbiota rhythmicity and infant CFI

Finally, we addressed whether gut microbiota rhythmicity relates to infant circadian maturation by testing interactions between zOTU sine and cosine fitting with CFI. Significant interactions would suggest that CFI relates to the rhythmic expression of bacterial composition. Indeed, this interaction was found in sine fitting for 27 zOTUs (1.81% of total zOTUs), with the number of rhythmic zOTUs linked to CFI increasing slightly across 3, 6 and 12 months to 9, 11 and 15 zOTUs, respectively (Figure 8A). We observed that the interaction of sine rhythmicity with CFI was mainly represented within *Actinobacteria* at 3 and 6 months (33.3% and 45.5% respectively, Figure 8B), and *Bacteroidetes* at 12 months (46.7%).

**Figure 8.**
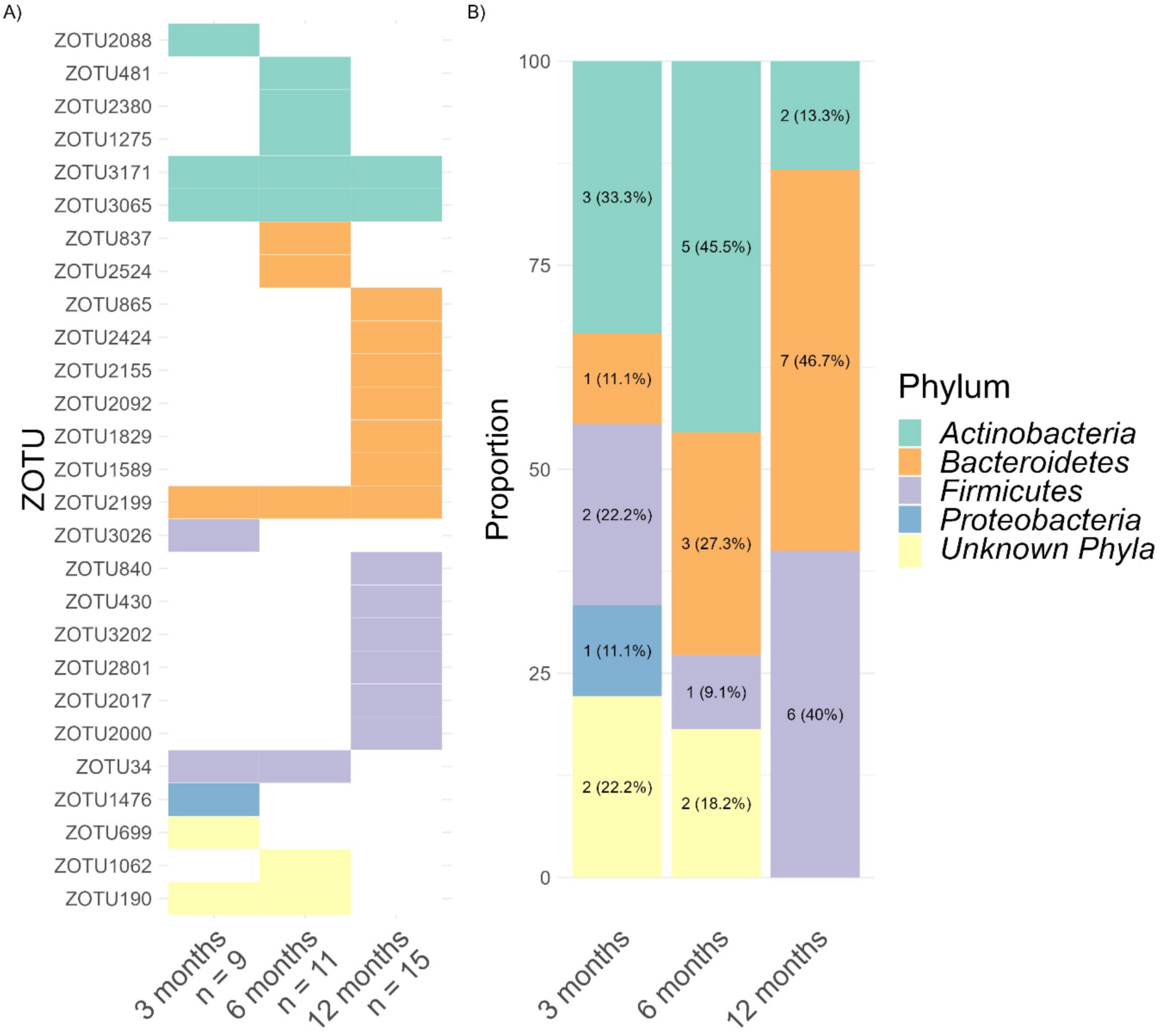
A) Heatmap depicting zOTUs (n = 27) with significantly associated sine rhythmicity and CFI (p < 0.05). The numbers below age groups indicate the number of zOTUs associated. B) Distribution of zOTUs phyla whose rhythmic sine patterns were linked with CFI.

Cosine microbial rhythms were associated with CFI in 96 zOTUs (corresponding to 6.43% of total zOTUs). At 3 months cosine rhythmicity in 9 zOTUs was linked to CFI, increasing to 13 zOTUs at 6 months, and 81 zOTUs at 12 months (Figure 9A), indicating an increasing number of zOTUs exhibiting cosine rhythmicity interacting with circadian rhythm maturation across the first year of life. At 3 months, most zOTUs linked to CFI were assigned to unknown phyla, while at 6 and 12 months mainly Firmicutes was involved (38.5% and 43.2%, Figure 9B).

**Figure 9.**
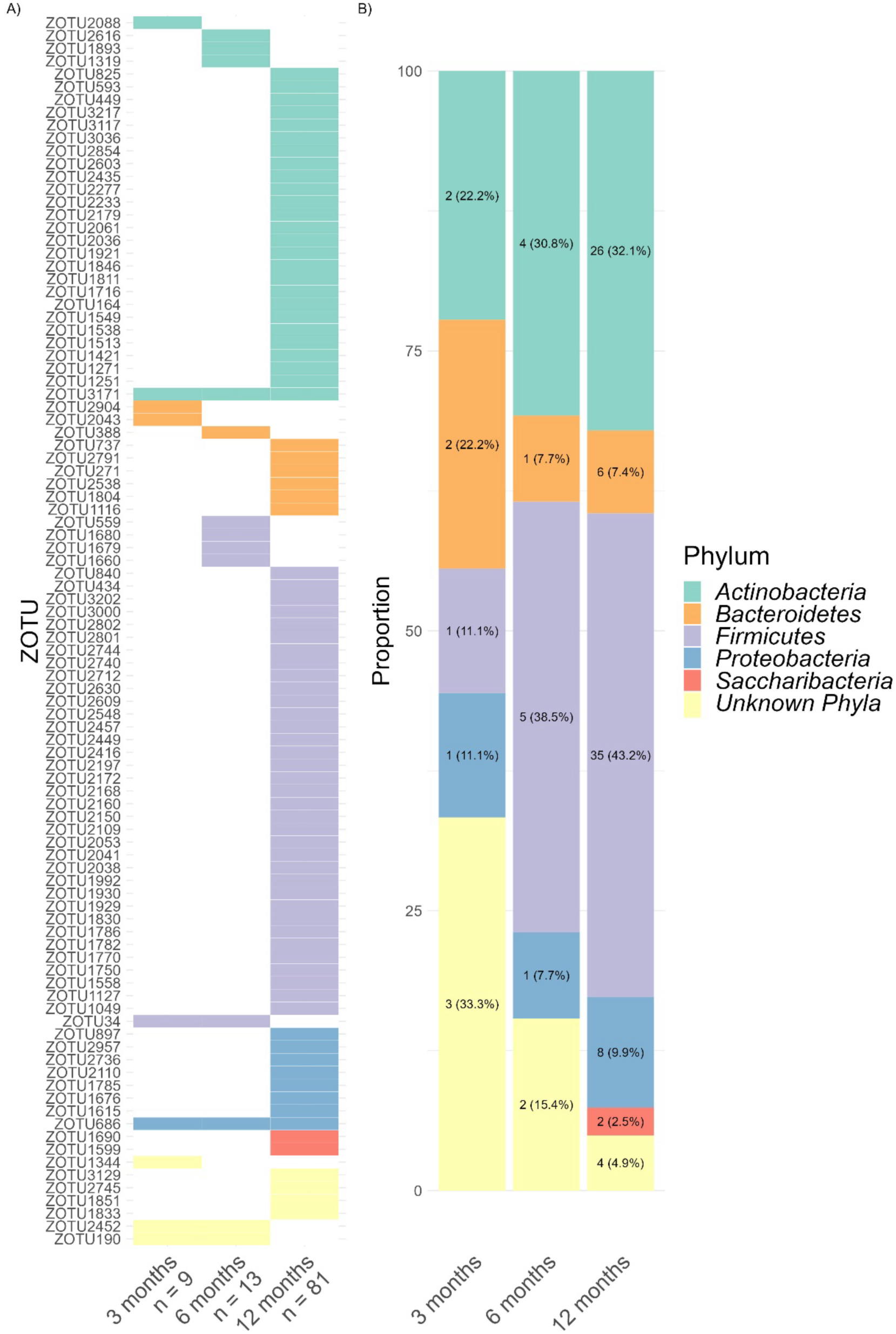
A) Heatmap indicating zOTUs (n = 96) with significantly associated cosine rhythm and CFI (p < 0.05). Values below age groups represent the number of corresponding zOTUs. B) Distribution of zOTUs phyla with cosine rhythmicity linked to CFI.

In summary, these results establish associations between circadian maturation in infants, quantified by the Circadian Function Index (CFI), and gut microbial taxa, involving 7.02% of zOTUs. Crucially, microbial composition exhibits diurnal variation, with 1.74% of zOTUs following a sine rhythm, and 6.69% adhering to a cosine rhythm. The expression of both, sine and cosine zOTU rhythmicity, gradually increases across infancy. Finally, the interaction of sine and cosine-fitted microbial rhythm with CFI was found in 1.81% of zOTUs for sine rhythm, and 6.43% of zOTUs for cosine rhythm, the latter experiencing a gradual strengthening with age.

## Discussion

In this longitudinal study across the first year of human life we examined the interaction between sleep rhythm maturation (CFI) and diurnal variation of gut microbiota. We demonstrated that CFI increases with age, confirming the measure as a reliable proxy for circadian maturation when using infant ankle actimetry. Interestingly, infant CFI was associated with gut microbial taxa, a link that strengthened with age. Specific Zero-radius Operational Taxonomic Units (zOTUs), primarily from the phyla *Actinobacteria*, *Firmicutes*, and *Bacteroidetes*, manifested rhythmic fluctuations across the day, following distinct sine or cosine patterns in their relative abundance. Age-related augmentation in rhythmicity was predominantly observed in cosine fitting, suggesting a gradual development of microbial diurnal rhythm with bacterial relative abundance peaking around midnight and reaching a nadir around midday. Remarkably, gut microbial rhythm was connected to CFI, underscoring a relationship between host circadian rhythm and gut microbial diurnality. Particularly cosine microbial rhythm maturation was accompanied with CFI. These findings demonstrate that sleep rhythm and gut diurnal profiles are associated in infancy and highlight the potential that infant sleep may interact with microbial dynamics, which may ultimately foster microbial colonization and developmental health.

This study employed CFI computed from ankle actimetry, which revealed a progressive increase in behavioral sleep rhythmicity. This finding aligns with developmental research indicating that sleep-wake patterns mature during the first year towards adult patterns (Zornoza-Moreno et al., 2011). The progressive increase in CFI from 3 to 12 months highlights a critical developmental window where sleep-wake patterns gradually evolve from fragmented to consolidated forms. This maturation is critical, as sleep quality, timing, and duration are linked to various aspects of developmental health, including cognitive functions like executive control, neural connectivity, and synaptic stability, all of which are crucial for neurodevelopment and cognitive maturation (Beaugrand et al., 2023; Kurth et al., 2012; LeBourgeois et al., 2019; Paavonen et al., 2010). Previous research on the development of sleep-wake patterns used actigraphy to objectively assess periods of sleep, capturing various metrics such as total sleep time, sleep efficiency, and wake after sleep onset (Schoch, Huber, et al., 2020). Our use of CFI provides a novel perspective on the maturation of infant sleep rhythms and supports CFI as a proxy for pediatric circadian rhythm research.

The findings confirmed the hypothesis that the circadian maturation during the first year of life is associated with gut microbiota composition. While both circadian maturation and the gut microbiome are expected developmental processes, our results provide novel insights into how these biological changes may be particularly linked. Specifically, we found that the number of bacterial taxa associated with CFI increases across infancy, more specifically showing that increased CFI is linked with higher relative abundance of *Bacteroidetes* and *Proteobacteria* and decreased abundance of *Firmicutes*. These results add to the evidence that circadian rhythm is associated with microbial colonization and the reciprocal interaction between gut microbiota rhythms and the host’s sleep pattern (Heppner et al., 2024; Thaiss et al., 2016). Our findings expand on this perspective by demonstrating that the relationship between gut microbiota and host rhythms encompasses not only sleep behavior but also the broader spectrum of activity-rest cycles extending beyond a single 24-hour cycle. This underscores the possibility of daily activity patterns in shaping gut microbiota composition, which likely impacts development. Conversely, bottom-up signals through gut microbiota may influence behavioral rhythmicity, which, in turn, likely affects long-term developmental outcomes (Carlson et al., 2018; Diaz Heijtz et al., 2011).

This study provides evidence for diurnal dynamics in the gut microbiota throughout infancy. Using sine and cosine fitting, we identified rhythmicity in specific bacterial taxa (zOTUs), with 26 showing rhythmicity in sine fitting and 100 in cosine fitting. As expected, data confirm that the rhythmicity of gut bacteria becomes more pronounced across increasing age groups, which aligns with recent research highlighting diurnal fluctuations in microbial and metabolic profiles in infants (Heppner et al., 2024). The increased rhythmicity of gut microbiota with age may be driven by several developmental changes. For instance, the transition from breastfeeding to solid foods significantly impacts gut microbiota composition, potentially increasing the abundance of rhythmic bacterial taxa (Bäckhed et al., 2015). Moreover, regular feeding schedules, which tend to become more consistent during the first year of life, are linked to greater microbial diversity, as shown in our previous research (Mühlematter et al., 2023). Feeding regularity may not only influence the composition of the gut microbiota but also support its rhythmicity, a relationship previously demonstrated in animal models (Thaiss et al., 2014). The effects of feeding rhythms on microbiota and host health may extend beyond infancy. In adults, studies have shown that meal timing plays a crucial role in aligning internal circadian rhythms, with benefits for metabolic health. For example, restricting meals to daytime has been found to prevent circadian misalignment and reduce glucose intolerance, a finding relevant for conditions like night shift work (Chellappa et al., 2021). Disruption of microbial rhythmicity has also been linked to metabolic health issues in adults, such as type 2 diabetes, suggesting that interventions supporting microbial rhythms could benefit long-term metabolic health (Reitmeier et al., 2020). Recognizing the importance of diurnal rhythms in gut microbiota could lead to new clinical and preventive strategies across the lifespan, especially in metabolic and immune health. Targeting microbial rhythmicity through dietary modifications or probiotic interventions may hold promise for supporting health outcomes from infancy through adulthood.

Our findings show that certain bacterial taxa in the gut microbiota exhibit daily rhythms in their relative abundance, which align with indicators of circadian maturation in infants. This suggests that microbial taxa gradually adjust their diurnal patterns, potentially coordinating with the infant’s sleep-wake and activity cycles. As infants’ sleep becomes more structured over the first year, certain microbial taxa accompany this host behavior, notably within the Bacteroidetes and Firmicutes phyla, which show increasing proportions as CFI scores rise. Bacteroidetes and Firmicutes are known contributors to SCFA production, including butyrate, acetate, and propionate, each of which has distinct roles in metabolic and neural pathways. For instance, butyrate, a major SCFA produced by Firmicutes (Vital et al., 2014), has been shown to influence sleep architecture in animal models by promoting non-rapid eye movement (NREM) sleep and affecting the regulation of neural circuits involved in sleep (Szentirmai et al., 2019). Studies in older adults suggest higher butyrate and propionate to be associated with improved sleep efficiency and longer sleep duration (Magzal et al., 2021). In the context of our findings, the increased relative abundance of SCFA-producing taxa (e.g., Firmicutes) in infants with higher CFI may indicate that microbial metabolic byproducts support sleep rhythm establishment. The gut-brain axis may mediate the gut microbiota communication with sleep and circadian rhythms, with the vagus nerve and intestinal epithelial cells transmitting microbial signals to the brain (Cryan et al., 2019; Eshleman & Alenghat, 2021; Valles-Colomer et al., 2019). Our findings suggest that higher SCFA-producing taxa, particularly within the Firmicutes and Bacteroidetes phyla, are linked to circadian maturation. Targeting these strains through dietary or probiotic interventions may offer a strategy to support sleep development, physiological resilience, and mental health during early life and beyond (Aburto & Cryan, 2024).

To our knowledge, no published studies have yet explicitly demonstrated age-related increases in CFI in infants, highlighting the need for further validation of this measure as a marker of circadian system maturation. To further elucidate human microbial rhythms and their health implications, future research should investigate environmental context, particularly diet, which is a key driver of gut microbiota composition (Weng et al., 2024). For example, high-fat diet is associated with increased Firmicutes and decreased Bacteroidetes, underscoring the effect of dietary patterns on microbial communities (Murphy et al., 2015).

Even though feeding categories were considered, our analysis lacks precise data on individual diet and the timing of solid food introduction. While our analysis used rigorous longitudinal actimetry data, the computation of microbacterial rhythms relies on cross-sectional sampling. Serial stool samples from the same infants, ideally including nighttime collections, would strengthen the assessment of microbial rhythmicity, as our current sampling was biased toward daytime and may have limited rhythm detection. The observed sine and cosine patterns represent trends across the observed period; but denser sampling could validate diurnal fits and uncover additional rhythmic cycles. Interactions between diet, microbial rhythms, and infant sleep patterns remain critical to understanding these complex developmental systems. Future research should identify specific bacterial strains and metabolic activity or products that promote host circadian function, paving the way for targeted pediatric therapies.

This study reveals rhythmic dynamics in gut microbiota across infancy and demonstrates the association between infant circadian rhythm and gut rhythmicity, emphasizing the importance of microbial rhythms in pediatric, neuromaturational, and immunological development.

Considering gut microbiota dynamics in early life could lead to targeted interventions for at-risk pediatric groups. For instance, dietary modifications aimed at promoting SCFA-producing bacteria or probiotic supplements containing specific bacterial strains associated with circadian support (e.g., certain Firmicutes or Bacteroidetes strains) could potentially enhance sleep regulation and overall developmental health. Tailoring early-life interventions in this way holds promise for managing health risks and supporting optimal growth trajectories through microbial health.

## Declarations

### Ethics approval and consent to participate

The cantonal ethics committee of Zurich, Switzerland, approved the project (BASEC 2016-00730), in line with the principles of the Helsinki Declaration. Before study enrollment, parents received detailed explanations about the study procedures before providing written consent.

## Funding

Swiss National Science Foundation (PCEFP1-181279 to SK; P0ZHP1-178697 to SS), University of Zurich (Medical Faculty; Forschungskredit FK-18-047, Clinical Research Priority Program “Sleep and Health”, to SK), Foundation for Research in Science and the Humanities (STWF-17-008, to SK), and Olga Mayenfisch Foundation (to SK).

### Availability of data

Data available on request due to privacy/ethical restrictions

### Declaration of AI-assisted technologies

During the preparation of this work the authors used chatGPT and perplexity to improve language. After using these AI tools, the authors reviewed and edited the content as needed and take full responsibility for the content of the publication.

### Authors’ contributions

CM and SK conceptualized the study. CM, SS, and JCM curated the data. Formal analysis was conducted by CM, SS, JCM, and JCW. SK and SS obtained funding. CM conducted the investigation. Methodology was developed by CM, SK, JCW, and DN. SK managed the project and provided resources together with SS. SK supervised the study. Validation was performed by CM, JCM, JCW, and SK. CM created visualizations. CM and SK wrote the original draft. All authors reviewed and edited the manuscript and approved the final version.

## Acknowledgments

We thank the participants for their dedication and the students and interns for their help with data collection.

## References

Aburto, M. R., & Cryan, J. F. (2024). Gastrointestinal and brain barriers: Unlocking gates of communication across the microbiota–gut–brain axis. Nature Reviews Gastroenterology & Hepatology, 21(4), 222–247. 10.1038/s41575-023-00890-0

Altaha, B., Heddes, M., Pilorz, V., Niu, Y., Gorbunova, E., Gigl, M., Kleigrewe, K., Oster, H., Haller, D., & Kiessling, S. (2022). Genetic and environmental circadian disruption induce weight gain through changes in the gut microbiome. Molecular Metabolism, 66, 101628. 10.1016/j.molmet.2022.101628

Bates, D., Mächler, M., Bolker, B., & Walker, S. (2015). Fitting Linear Mixed-Effects Models Using lme4. Journal of Statistical Software, 67, 1–48. 10.18637/jss.v067.i01

Beaugrand, M., Muehlematter, C., Markovic, A., Camos, V., & Kurth, S. (2023). Sleep as a protective factor of children’s executive functions: A study during COVID-19 confinement. PLOS ONE, 18(1), e0279034. 10.1371/journal.pone.0279034

Besedovsky, L., Lange, T., & Haack, M. (2019). The Sleep-Immune Crosstalk in Health and Disease. Physiological Reviews, 99(3), 1325–1380. 10.1152/physrev.00010.2018

Blume, C., Santhi, N., & Schabus, M. (2016). ‘nparACT’ package for R: A free software tool for the non-parametric analysis of actigraphy data. MethodsX, 3, 430–435. 10.1016/j.mex.2016.05.006

Bowers, S. J., Vargas, F., González, A., He, S., Jiang, P., Dorrestein, P. C., Knight, R., Jr, K. P. W., Lowry, C. A., Fleshner, M., Vitaterna, M. H., & Turek, F. W. (2020). Repeated sleep disruption in mice leads to persistent shifts in the fecal microbiome and metabolome. PLOS ONE, 15(2), e0229001. 10.1371/journal.pone.0229001

Brown, R., Price, R. J., King, M. G., & Husband, A. J. (1990). Are antibiotic effects on sleep behavior in the rat due to modulation of gut bacteria? Physiology & Behavior, 48(4), 561–565. 10.1016/0031-9384(90)90300-S

Carlson, A. L., Xia, K., Azcarate-Peril, M. A., Goldman, B. D., Ahn, M., Styner, M. A., Thompson, A. L., Geng, X., Gilmore, J. H., & Knickmeyer, R. C. (2018). Infant Gut Microbiome Associated With Cognitive Development. Biological Psychiatry, 83(2), 148–159. 10.1016/j.biopsych.2017.06.021

Carpena, M. X., Barros, A. JD., Comelli, E. M., López-Domínguez, L., Alves, E. D., Wendt, A., Crochemore-Silva, I., Bandsma, R. HJ., Santos, I. S., Matijasevich, A., Borges, M. C., & Tovo-Rodrigues, L. (2024). Accelerometer-based sleep metrics and gut microbiota during adolescence: Association findings from a Brazilian population-based birth cohort. Sleep Medicine, 114, 203–209. 10.1016/j.sleep.2023.12.028

Catassi, G., Aloi, M., Giorgio, V., Gasbarrini, A., Cammarota, G., & Ianiro, G. (2024). The Role of Diet and Nutritional Interventions for the Infant Gut Microbiome. Nutrients, 16(3), 400. 10.3390/nu16030400

Chellappa, S. L., Qian, J., Vujovic, N., Morris, C. J., Nedeltcheva, A., Nguyen, H., Rahman, N., Heng, S. W., Kelly, L., Kerlin-Monteiro, K., Srivastav, S., Wang, W., Aeschbach, D., Czeisler, C. A., Shea, S. A., Adler, G. K., Garaulet, M., & Scheer, F. A. J. L. (2021). Daytime eating prevents internal circadian misalignment and glucose intolerance in night work. Science Advances, 7(49), eabg9910. 10.1126/sciadv.abg9910

Cornelissen, G. (2014). Cosinor-based rhythmometry. Theoretical Biology & Medical Modelling, 11, 16. 10.1186/1742-4682-11-16

Cryan, J. F., O’Riordan, K. J., Cowan, C. S. M., Sandhu, K. V., Bastiaanssen, T. F. S., Boehme, M., Codagnone, M. G., Cussotto, S., Fulling, C., Golubeva, A. V., Guzzetta, K. E., Jaggar, M., Long-Smith, C. M., Lyte, J. M., Martin, J. A., Molinero-Perez, A., Moloney, G., Morelli, E., Morillas, E.,… Dinan, T. G. (2019). The Microbiota-Gut-Brain Axis. Physiological Reviews, 99(4), 1877–2013. 10.1152/physrev.00018.2018

DeSantis, T. Z., Hugenholtz, P., Larsen, N., Rojas, M., Brodie, E. L., Keller, K., Huber, T., Dalevi, D., Hu, P., & Andersen, G. L. (2006). Greengenes, a chimera-checked 16S rRNA gene database and workbench compatible with ARB. Applied and Environmental Microbiology, 72(7), 5069–5072. 10.1128/AEM.03006-05

Diaz Heijtz, R., Wang, S., Anuar, F., Qian, Y., Björkholm, B., Samuelsson, A., Hibberd, M. L., Forssberg, H., & Pettersson, S. (2011). Normal gut microbiota modulates brain development and behavior. Proceedings of the National Academy of Sciences of the United States of America, 108(7), 3047–3052. 10.1073/pnas.1010529108

Edgar, R. C., & Flyvbjerg, H. (2015). Error filtering, pair assembly and error correction for next-generation sequencing reads. *Bioinformatics (Oxford*, England), 31(21), 3476– 3482. 10.1093/bioinformatics/btv401

Eshleman, E. M., & Alenghat, T. (2021). Epithelial sensing of microbiota-derived signals. Genes & Immunity, 22(5), 237–246. 10.1038/s41435-021-00124-w

Grosicki, G. J., Riemann, B. L., Flatt, A. A., Valentino, T., & Lustgarten, M. S. (2020). Self-reported sleep quality is associated with gut microbiome composition in young, healthy individuals: A pilot study. Sleep Medicine, 73, 76–81. 10.1016/j.sleep.2020.04.013

Heddes, M., Altaha, B., Niu, Y., Reitmeier, S., Kleigrewe, K., Haller, D., & Kiessling, S. (2022). The intestinal clock drives the microbiome to maintain gastrointestinal homeostasis. Nature Communications, 13(1), 6068. 10.1038/s41467-022-33609-x

Heppner, N., Reitmeier, S., Heddes, M., Merino, M. V., Schwartz, L., Dietrich, A., List, M., Gigl, M., Meng, C., Veen, D. R. van der, Schirmer, M., Kleigrewe, K., Omer, H., Kiessling, S., & Haller, D. (2024). Diurnal rhythmicity of infant fecal microbiota and metabolites: A randomized controlled interventional trial with infant formula. Cell Host & Microbe, 32(4), 573–587.e5. 10.1016/j.chom.2024.02.015

Hill, C. J., Lynch, D. B., Murphy, K., Ulaszewska, M., Jeffery, I. B., O’Shea, C. A., Watkins, C., Dempsey, E., Mattivi, F., Touhy, K., Ross, R. P., Ryan, C. A., O’ Toole, P. W., & Stanton, C. (2017). Evolution of gut microbiota composition from birth to 24 weeks in the INFANTMET Cohort. Microbiome, 5. 10.1186/s40168-016-0213-y

Iglowstein, I., Jenni, O. G., Molinari, L., & Largo, R. H. (2003). Sleep Duration From Infancy to Adolescence: Reference Values and Generational Trends. Pediatrics, 111(2), 302–307. 10.1542/peds.111.2.302

Jaramillo, V., Schoch, S. F., Markovic, A., Kohler, M., Huber, R., Lustenberger, C., & Kurth, S. (2023). An infant sleep electroencephalographic marker of thalamocortical connectivity predicts behavioral outcome in late infancy. NeuroImage, 269, 119924. 10.1016/j.neuroimage.2023.119924

Korostovtseva, L., Bochkarev, M., & Sviryaev, Y. (2021). Sleep and Cardiovascular Risk. Sleep Medicine Clinics, 16(3), 485–497. 10.1016/j.jsmc.2021.05.001

Krych, Ł., Kot, W., Bendtsen, K. M. B., Hansen, A. K., Vogensen, F. K., & Nielsen, D. S. (2018). Have you tried spermine? A rapid and cost-effective method to eliminate dextran sodium sulfate inhibition of PCR and RT-PCR. Journal of Microbiological Methods, 144, 1–7. 10.1016/j.mimet.2017.10.015

Kurth, S., Ringli, M., Lebourgeois, M. K., Geiger, A., Buchmann, A., Jenni, O. G., & Huber, R. (2012). Mapping the electrophysiological marker of sleep depth reveals skill maturation in children and adolescents. NeuroImage, 63(2), 959–965. 10.1016/j.neuroimage.2012.03.053

LeBourgeois, M. K., Dean, D. C., Deoni, S. C. L., Kohler, M., & Kurth, S. (2019). A simple sleep EEG marker in childhood predicts brain myelin 3.5 years later. NeuroImage, 199, 342–350. 10.1016/j.neuroimage.2019.05.072

Leiner, D. J. (2021). *SoSci Survey (Version 3.2.31) [Computer software]* [Computer software]. Available at https://www.soscisurvey.de

Lubin, J.-B., Green, J., Maddux, S., Denu, L., Duranova, T., Lanza, M., Wynosky-Dolfi, M., Flores, J. N., Grimes, L. P., Brodsky, I. E., Planet, P. J., & Silverman, M. A. (2023). Arresting microbiome development limits immune system maturation and resistance to infection in mice. Cell Host & Microbe, 31(4), 554–570.e7. 10.1016/j.chom.2023.03.006

Magoč, T., & Salzberg, S. L. (2011). FLASH: Fast length adjustment of short reads to improve genome assemblies. Bioinformatics, 27(21), 2957–2963. 10.1093/bioinformatics/btr507

Magzal, F., Even, C., Haimov, I., Agmon, M., Asraf, K., Shochat, T., & Tamir, S. (2021). Associations between fecal short-chain fatty acids and sleep continuity in older adults with insomnia symptoms. Scientific Reports, 11(1), 4052. 10.1038/s41598-021-83389-5

Martin, M. (2011). Cutadapt removes adapter sequences from high-throughput sequencing reads. EMBnet.Journal, 17(1), Article 1. 10.14806/ej.17.1.200

McMurdie, P. J., & Holmes, S. (2013). phyloseq: An R Package for Reproducible Interactive Analysis and Graphics of Microbiome Census Data. PLOS ONE, 8(4), e61217. 10.1371/journal.pone.0061217

Mercer, E. M., Ramay, H. R., Moossavi, S., Laforest-Lapointe, I., Reyna, M. E., Becker, A. B., Simons, E., Mandhane, P. J., Turvey, S. E., Moraes, T. J., Sears, M. R., Subbarao, P., Azad, M. B., & Arrieta, M.-C. (2024). Divergent maturational patterns of the infant bacterial and fungal gut microbiome in the first year of life are associated with inter-kingdom community dynamics and infant nutrition. Microbiome, 12(1), 22. 10.1186/s40168-023-01735-3

Mühlematter, C., Nielsen, D. S., Castro-Mejía, J. L., Brown, S. A., Rasch, B., Jr, K. P. W., Walser, J.-C., Schoch, S. F., & Kurth, S. (2023). Not simply a matter of parents— Infants’ sleep-wake patterns are associated with their regularity of eating. PLOS ONE, 18(10), e0291441. 10.1371/journal.pone.0291441

Mukherji, A., Kobiita, A., Ye, T., & Chambon, P. (2013). Homeostasis in Intestinal Epithelium Is Orchestrated by the Circadian Clock and Microbiota Cues Transduced by TLRs. Cell, 153(4), 812–827. 10.1016/j.cell.2013.04.020

Murphy, E. A., Velazquez, K. T., & Herbert, K. M. (2015). Influence of high-fat diet on gut microbiota: A driving force for chronic disease risk. Current Opinion in Clinical Nutrition & Metabolic Care, 18(5), 515. 10.1097/MCO.0000000000000209

Nonaka, K., Nakazawa, Y., & Kotorii, T. (1983). Effects of antibiotics, minocycline and ampicillin, on human sleep. Brain Research, 288(1–2), 253–259. 10.1016/0006-8993(83)90101-4

Ogawa, Y., Miyoshi, C., Obana, N., Yajima, K., Hotta-Hirashima, N., Ikkyu, A., Kanno, S., Soga, T., Fukuda, S., & Yanagisawa, M. (2020). Gut microbiota depletion by chronic antibiotic treatment alters the sleep/wake architecture and sleep EEG power spectra in mice. Scientific Reports, 10(1), Article 1. 10.1038/s41598-020-76562-9

Ortiz-Tudela, E., Martinez-Nicolas, A., Campos, M., Rol, M. Á., & Madrid, J. A. (2010). A New Integrated Variable Based on Thermometry, Actimetry and Body Position (TAP) to Evaluate Circadian System Status in Humans. PLOS Computational Biology, 6(11), e1000996. 10.1371/journal.pcbi.1000996

Ovreås, L., Forney, L., Daae, F. L., & Torsvik, V. (1997). Distribution of bacterioplankton in meromictic Lake Saelenvannet, as determined by denaturing gradient gel electrophoresis of PCR-amplified gene fragments coding for 16S rRNA. Applied and Environmental Microbiology. https://journals.asm.org/doi/abs/10.1128/aem.63.9.3367-3373.1997

Paavonen, E. J., Räikkönen, K., Pesonen, A.-K., Lahti, J., Komsi, N., Heinonen, K., Järvenpää, A.-L., Strandberg, T., Kajantie, E., & Porkka-Heiskanen, T. (2010). Sleep quality and cognitive performance in 8-year-old children. Sleep Medicine, 11(4), 386–392. 10.1016/j.sleep.2009.09.009

Pantazi, A. C., Balasa, A. L., Mihai, C. M., Chisnoiu, T., Lupu, V. V., Kassim, M. A. K., Mihai, L., Frecus, C. E., Chirila, S. I., Lupu, A., Andrusca, A., Ionescu, C., Cuzic, V., & Cambrea, S. C. (2023). Development of Gut Microbiota in the First 1000 Days after Birth and Potential Interventions. Nutrients, 15(16). 10.3390/nu15163647

Reitmeier, S., Kiessling, S., Clavel, T., List, M., Almeida, E. L., Ghosh, T. S., Neuhaus, K., Grallert, H., Linseisen, J., Skurk, T., Brandl, B., Breuninger, T. A., Troll, M., Rathmann, W., Linkohr, B., Hauner, H., Laudes, M., Franke, A., Le Roy, C. I.,… Haller, D. (2020). Arrhythmic Gut Microbiome Signatures Predict Risk of Type 2 Diabetes. Cell Host & Microbe, 28(2), 258–272.e6. 10.1016/j.chom.2020.06.004

Schmieder, R., & Edwards, R. (2011). Quality control and preprocessing of metagenomic datasets. Bioinformatics, 27(6), 863–864. 10.1093/bioinformatics/btr026

Schoch, S. F., Castro-Mejía, J. L., Krych, L., Leng, B., Kot, W., Kohler, M., Huber, R., Rogler, G., Biedermann, L., Walser, J. C., Nielsen, D. S., & Kurth, S. (2022). From Alpha Diversity to Zzz: Interactions among sleep, the brain, and gut microbiota in the first year of life. Progress in Neurobiology, 209, 102208. 10.1016/j.pneurobio.2021.102208

Schoch, S. F., Huber, R., Kohler, M., & Kurth, S. (2020). Which Are the Central Aspects of Infant Sleep? The Dynamics of Sleep Composites across Infancy. Sensors, 20(24), Article 24. 10.3390/s20247188

Schoch, S. F., Jenni, O. G., Kohler, M., & Kurth, S. (2019). Actimetry in infant sleep research: An approach to facilitate comparability. Sleep, 42(7), zsz083. 10.1093/sleep/zsz083

Schoch, S. F., Kurth, S., & Werner, H. (2020). Actigraphy in sleep research with infants and young children: Current practices and future benefits of standardized reporting. *Journal of Sleep Research*, e13134. 10.1111/jsr.13134

Sen, P., Molinero-Perez, A., O’Riordan, K. J., McCafferty, C. P., O’Halloran, K. D., & Cryan, J. F. (2021). Microbiota and sleep: Awakening the gut feeling. Trends in Molecular Medicine, 27(10), 935–945. 10.1016/j.molmed.2021.07.004

Sinthong, A., & Ngernlangtawee, D. (2024). Early sleep intervention for improving infant sleep quality: A randomized controlled trial, preliminary result. BMC Pediatrics, 24(1), 306. 10.1186/s12887-024-04771-6

Smith, R. P., Easson, C., Lyle, S. M., Kapoor, R., Donnelly, C. P., Davidson, E. J., Parikh, E., Lopez, J. V., & Tartar, J. L. (2019). Gut microbiome diversity is associated with sleep physiology in humans. PLOS ONE, 14(10), e0222394. 10.1371/journal.pone.0222394

Sordillo, J. E., Korrick, S., Laranjo, N., Carey, V., Weinstock, G. M., Gold, D. R., O’Connor, G., Sandel, M., Bacharier, L. B., Beigelman, A., Zeiger, R., Litonjua, A. A., & Weiss, S. T. (2019). Association of the Infant Gut Microbiome With Early Childhood Neurodevelopmental Outcomes: An Ancillary Study to the VDAART Randomized Clinical Trial. JAMA Network Open, 2(3), e190905. 10.1001/jamanetworkopen.2019.0905

Stich, F. M., Huwiler, S., D’Hulst, G., & Lustenberger, C. (2022). The Potential Role of Sleep in Promoting a Healthy Body Composition: Underlying Mechanisms Determining Muscle, Fat, and Bone Mass and Their Association with Sleep. Neuroendocrinology, 112(7), 673–701. 10.1159/000518691

Szentirmai, É., Millican, N. S., Massie, A. R., & Kapás, L. (2019). Butyrate, a metabolite of intestinal bacteria, enhances sleep. Scientific Reports, 9(1), 1–9. 10.1038/s41598-019-43502-1

Tahara, Y., Yamazaki, M., Sukigara, H., Motohashi, H., Sasaki, H., Miyakawa, H., Haraguchi, A., Ikeda, Y., Fukuda, S., & Shibata, S. (2018). Gut Microbiota-Derived Short Chain Fatty Acids Induce Circadian Clock Entrainment in Mouse Peripheral Tissue. Scientific Reports, 8(1), 1395. 10.1038/s41598-018-19836-7

Thaiss, C. A., Levy, M., Korem, T., Dohnalová, L., Shapiro, H., Jaitin, D. A., David, E., Winter, D. R., Gury-BenAri, M., Tatirovsky, E., Tuganbaev, T., Federici, S., Zmora, N., Zeevi, D., Dori-Bachash, M., Pevsner-Fischer, M., Kartvelishvily, E., Brandis, A., Harmelin, A.,… Elinav, E. (2016). Microbiota Diurnal Rhythmicity Programs Host Transcriptome Oscillations. Cell, 167(6), 1495–1510.e12. 10.1016/j.cell.2016.11.003

Thaiss, C. A., Zeevi, D., Levy, M., Zilberman-Schapira, G., Suez, J., Tengeler, A. C., Abramson, L., Katz, M. N., Korem, T., Zmora, N., Kuperman, Y., Biton, I., Gilad, S., Harmelin, A., Shapiro, H., Halpern, Z., Segal, E., & Elinav, E. (2014). Transkingdom Control of Microbiota Diurnal Oscillations Promotes Metabolic Homeostasis. Cell, 159(3), 514–529. 10.1016/j.cell.2014.09.048

Timofeev, I., Schoch, S. F., LeBourgeois, M. K., Huber, R., Riedner, B. A., & Kurth, S. (2020). Spatio-temporal properties of sleep slow waves and implications for development. Current Opinion in Physiology, 15, 172–182. 10.1016/j.cophys.2020.01.007

Valles-Colomer, M., Falony, G., Darzi, Y., Tigchelaar, E. F., Wang, J., Tito, R. Y., Schiweck, C., Kurilshikov, A., Joossens, M., Wijmenga, C., Claes, S., Van Oudenhove, L., Zhernakova, A., Vieira-Silva, S., & Raes, J. (2019). The neuroactive potential of the human gut microbiota in quality of life and depression. Nature Microbiology, 4(4), 623–632. 10.1038/s41564-018-0337-x

Vital, M., Howe, A. C., & Tiedje, J. M. (2014). Revealing the Bacterial Butyrate Synthesis Pathways by Analyzing (Meta)genomic Data. mBio, 5(2), 10.1128/mbio.00889-14

Voigt, R. M., Forsyth, C. B., Green, S. J., Mutlu, E., Engen, P., Vitaterna, M. H., Turek, F. W., & Keshavarzian, A. (2014). Circadian Disorganization Alters Intestinal Microbiota. PLoS ONE, 9(5), e97500. 10.1371/journal.pone.0097500

Wang, Y., van de Wouw, M., Drogos, L., Vaghef-Mehrabani, E., Reimer, R. A., Tomfohr-Madsen, L., & Giesbrecht, G. F. (2022). Sleep and the gut microbiota in preschool-aged children. *Sleep*, zsac020. 10.1093/sleep/zsac020

Weng, H., Deng, L., Wang, T., Xu, H., Wu, J., Zhou, Q., Yu, L., Chen, B., Huang, L., Qu, Y., Zhou, L., & Chen, X. (2024). Humid heat environment causes anxiety-like disorder via impairing gut microbiota and bile acid metabolism in mice. Nature Communications, 15(1), 5697. 10.1038/s41467-024-49972-w

Werner, H., Molinari, L., Guyer, C., & Jenni, O. G. (2008). Agreement Rates Between Actigraphy, Diary, and Questionnaire for Children’s Sleep Patterns. Archives of Pediatrics & Adolescent Medicine, 162(4), 350–358. 10.1001/archpedi.162.4.350

Wickham, H. (2016). ggplot2: Elegant Graphics for Data Analysis (Springer-Verlag New York). https://ggplot2.tidyverse.org

Xiang, X., Chen, J., Zhu, M., Gao, H., Liu, X., & Wang, Q. (2023). Multiomics Revealed the Multi-Dimensional Effects of Late Sleep on Gut Microbiota and Metabolites in Children in Northwest China. Nutrients, 15(20), Article 20. 10.3390/nu15204315

Zornoza-Moreno, M., Fuentes-Hernández, S., Sánchez-Solis, M., Rol, M. Á., Larqué, E., & Madrid, and J. A. (2011). Assessment of Circadian Rhythms of Both Skin Temperature and Motor Activity in Infants During the First 6 Months of Life. Chronobiology International, 28(4), 330–337. 10.3109/07420528.2011.565895

